# Expressing of Cytochrome-c, ADAM 17, Wnt-5a, and Hedgehog gene during the tissue regeneration of digit tip mice (*Mus musculus*) var Swiss Webster post amputation

**DOI:** 10.1101/833491

**Authors:** Titta Novianti, Febriana Dwi Wahyuni, It Jamilah, Syafruddin Ilyas

**Affiliations:** Program Studi Bioteknologi, Universitas Esa Unggul, Jakarta Jl. Raya Arjuna Utara no. 9 Jakarta Barat; Program Studi Biologi, Universitas Sumatera Utara, Medan, Jl. Abdul Hakim No.1, Padang Bulan, Kec. Medan Baru, Kota Medan, Sumatera Utara 20222

**Keywords:** digit tip mice, ADAM 17, Cyt-c, Wnt-5a, Hedgehog

## Abstract

The tissue regeneration of digit tip mice needs some proteins that play a role in overcoming the inflammatory state. The functional protein plays a role in the continuous growth of specific cells, continuous migration, functional differentiation, and tissue morphogenesis. All of the cells need energy related to cell respiration. Naturally expressing mRNA of ADAM 17, Wnt-5a, Hedgehog (HH), and Cytochrome-c (Cyt-c) reliably produced the accordance with their respective roles in each specific phase of tissue regeneration until the whole tissues formed again. The ADAM 17 gen expressed in the inflammatory phase, it positively related its essential role to the inflammatory process. Cyt-c gene expression naturally occurs throughout the tissue regeneration because of its key role in the cellular respiration. Expressed Wnt-5a gene mRNA in the granulation phase, the specific HH gene expressed after the blastema phase. Both expressed genes positively correlate with the continual growth of the digit tip mice by the specific Spearman test (p <0.05) because of their active role of cell proliferation, cell differentiation, extensive migration, and morphogenesis.

## Background

Tissue regeneration is natural that will occur when a specific organism injured or amputated (Reinke and Sorg, 2012). Not all organisms typically undergo complete regeneration. Organisms that have naturally limited regeneration ability, possible injuries that naturally occur will merely be covered in scar tissue but cannot sufficiently restore lost organs if amputated. The lower animal levels, the higher the unique ability of extensive regeneration. Hydra, tapeworms, starfish, naturally have a high ability regeneration when neglecting parts of his specific body (Bedelbaeva et al., 2010; Tahara et al., 2018).

Vertebrate animals naturally have a limited ability in the extensive regeneration of specific organs and tissue. The ability fish in extensive regeneration in their prominent fins, urodella in its distinctive tail, modern lizards and geckos can regenerate naturally their distinctive tails after autotomy (Novianti et al., 2019; Fisher et al., 2012). Considered humans, as model organisms with the most superior levels of distinct taxa, limited ability in extensive regeneration. Therefore, a published study about tissue regeneration continues thoughtfully to develop naturally as a therapeutic effort is likely humans when injured or amputated (Weidemann and Johnson, 2008; Guedelhoefer and Alvarado, 2012).

Tissue regeneration naturally involves various specific cells, specific molecules, specific proteins, and specific genes. This complicated enough and complex due to specific tissue in managing all instruments and private compartments (Reinke and Sorg, 2012). There in common are four distinct stages of tissue regeneration, the wound-healing phase, the blastema phase, the regeneration phase, and the completed last in common is the maturation. The wound healing phase naturally occurs after the specific tissue has intentionally injured. In this sufficiently completed the completed phase, the specific tissue will typically experience local inflammation, granulation, and wound contraction. In each completed phase, adequately expressing specific genes, specific proteins, specific molecules, and specific cells typically involved in tissue regeneration is different. In the inflammatory phase, the specific tissue typically dominated white blood cells whose key role in phagocytosis the damaged cells. In the inflammatory phase, the specific tissue typically dominated white blood cells whose key role in phagocytosis the damaged cells (Mescher, 2017).

The following stage of the wound healing phase is the contraction of the extensive wound marked by the typically migrating of endothelial cells. At this distinct stage, fibroblast and macrophage cells in common were merely extensive migration and proliferation. The fibroblast cells will carefully secrete growth factors that will naturally stimulate forming the extracellular matrix. Macrophage cells play role cleansing tissue from the frequent rest of the extracellular matrix, so there is an appropriate balance between naturally producing and gradual degradation of the extracellular matrix (Krafts, 2010; Wynn and Vannella, 2016).

Tissue regeneration universally requires possible energy, so the dynamic process of cellular respiration in the mitochondria will progressively increase (Jornayvaz and Shulman, 2010). Cytochrome-c (Cyt-c) obtain a peripheral protein in the inner membrane of mitochondria. Cyt-c is synthesized in the cytosol and translocated to the mitochondria. The typically composed of Cyt-c in common is a single polypeptide chain of 104 amino acid residues. The specific functions of Cyt-c in the respiratory chain as an electron shuttle between complex III and complex IV and inhibits reactive oxygen species (ROS) formation, therefore prevents oxidative stress. The higher the cellular respiration process, the higher the Cyt-c expressed. The possible disruption of Cyt-c gene causes embryonic lethality, loosely and tightly bound of possible Cyt-c pools have been implicated in various functions of the specific organ (Wright et al., 2007; Panigrahy et al., 2013).

In the inflammatory phase, some leukocytes regulated the proteolytic processes that it was a significant role in modulating inflammation (Vitulo et al., 2017). The proteins on the surface of leukocyte cells lead to the release of a soluble extracellular domain fragment in the inflammatory phase. One of these proteins is a disintegrin and metalloproteinase-17 (ADAM17) that have a role in mediating the release of a soluble extracellular domain fragment. The role of ADAM17 in modulating inflammation process is not clear, but deficiency of leukocyte-expressed ADAM17- null mice in all leukocytes implicated in acute lung inflammation (Wang et al., 2019; Chalaris et al., 2010).

During the tissue regeneration process, the specific tissue was typically dominated by the proliferated, differentiated, and migrated cells. The complex process naturally required some specific protein that plays a role in tissue regeneration (Wynn and Vannella, 2016). Hedgehog (HH) protein plays an essential role in ion programs and extensive cell division required. In the mature adult, specific HH protein continues thoughtfully to discrete of the stem cell and progenitor cells within various organs, including the intact skin, brain prostate, and bladder. The specific functions of specific HH protein are accurately controlling the proliferation, functional specification and plasticity cells, whether the effective mechanism remains unclear (Petrova and Joyner, 2014; Lozito and Tuan, 2015).

Specific Wnt-5a genes encode the signaling molecules that have a role in regulating cell fate, adhesion, shape, proliferation, functional differentiation, and active movement, and naturally required for the possible development of multiple organs. Wnt-5a has naturally required for the proliferation of limb bud and progenitor cells, and it has contributed positively to the immune responses. Specific Wnt-5a protein properly regulating the functional differentiation of specialized T cells, naturally inducing IL-6 and active IL-1b expression, and necessary maintenance of innate immune responses. WNT-5a is equally critical in the effective regulation of functional differentiation and lineage commitment of the mesenchymal stem cell. The depletion of WNT-5a in MSCs leads to considerable loss of osteocyte producing capacity (Rishikaysh et al., 2014; Vitulo et al., 2017).

In this study, the analysis of Cyt-c gene expression was assumed to have a role in the process of tissue regeneration because of its role in the cell respiration process (Allen, 2011). Comparative analysis of ADAM 17 gene expression was correctly predicted it related to the inflammatory process, and analysis of the Wnt-5a and HH gene expression that play a role in the specific process of cell proliferation, functional differentiation, extensive migration, and morphogenesis (Wang et al., 2019; Petrova and Joyner, 2014; Kumawat and Gosens, 2016). We suggest there is the dynamic of some genes expression that each expression is various for every phase during the tissue regeneration process. In this comparative study, we naturally used the growth tissue from digit tip mice (*Mus musculus*) that were amputated.

## Results

### The Growth of digit tip mice (*Mus musculus*)

The continuous growth of digit tip mice (*Mus musculus*) after amputation (fig. 1) from day 0 (4 hours after amputation) until day 25. On day 0 and day 1 there is no visible growth of specific tissue, it was believed that the inflamed tissue. On day 3 until day 10, the growth tissue typically appeared, that the naturally formed of cell division but has not yet formed the new tissue. On day 15 and day 25, the new tissue has formed and formed the current nails.

**Figure 1.**
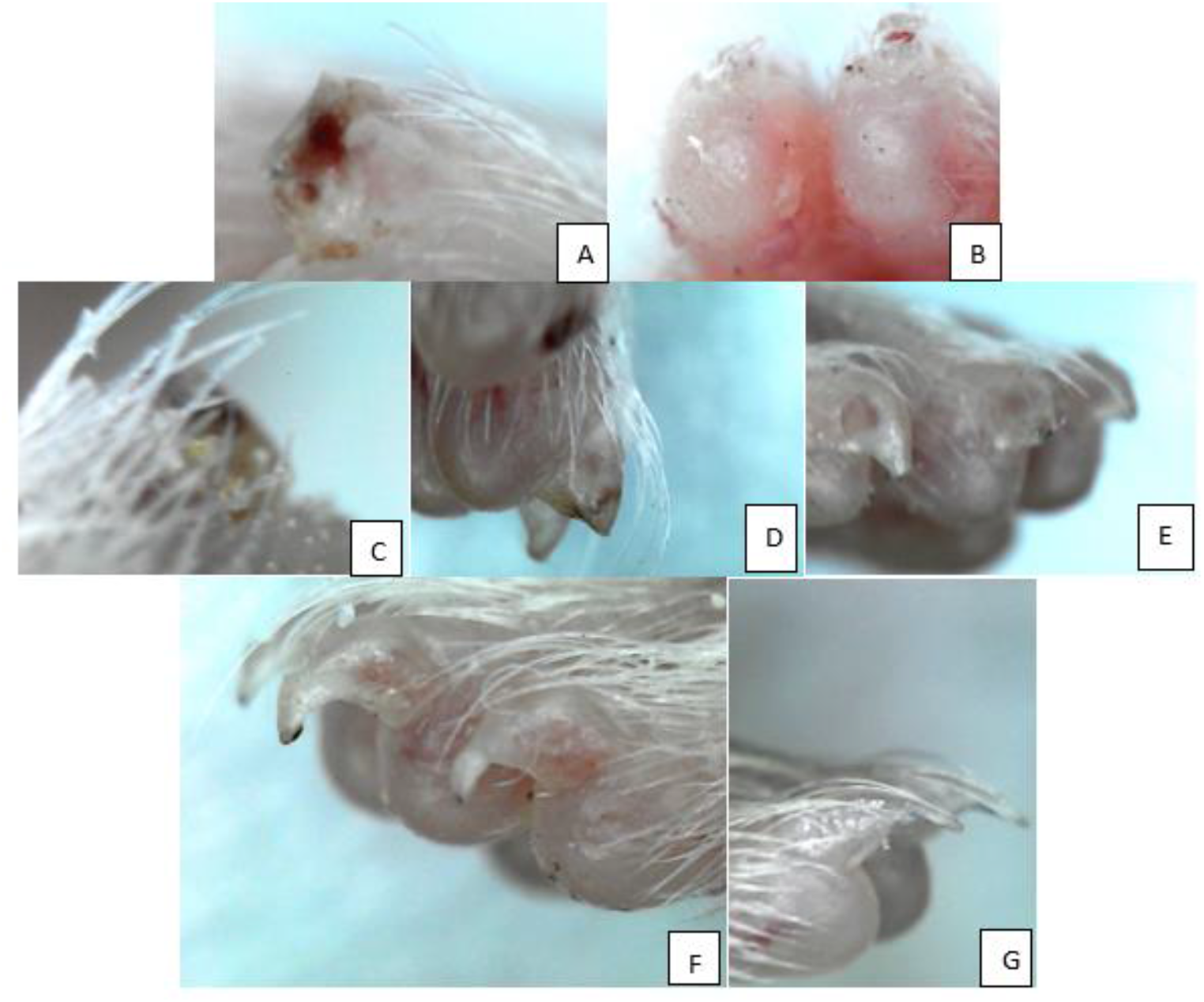
Growth of digit tip mice (*Mus musculus*) until day 25. A. The digit tip on day 0 (4 hours after amputation); B. Day 1 after amputation; C. Day 3 after amputation; D. Day 5 after amputation; E. Day 15 after amputation; F. Day 25 after amputation; G. Control

The curve of tissue growth of digit tip mice was shown in Fig 2. The curve grows slowly on day 0 (4 hours after amputation) until day 10 after amputation. After day 10, the curve line appears to increase sharply until day 25.

**Figure 2.**
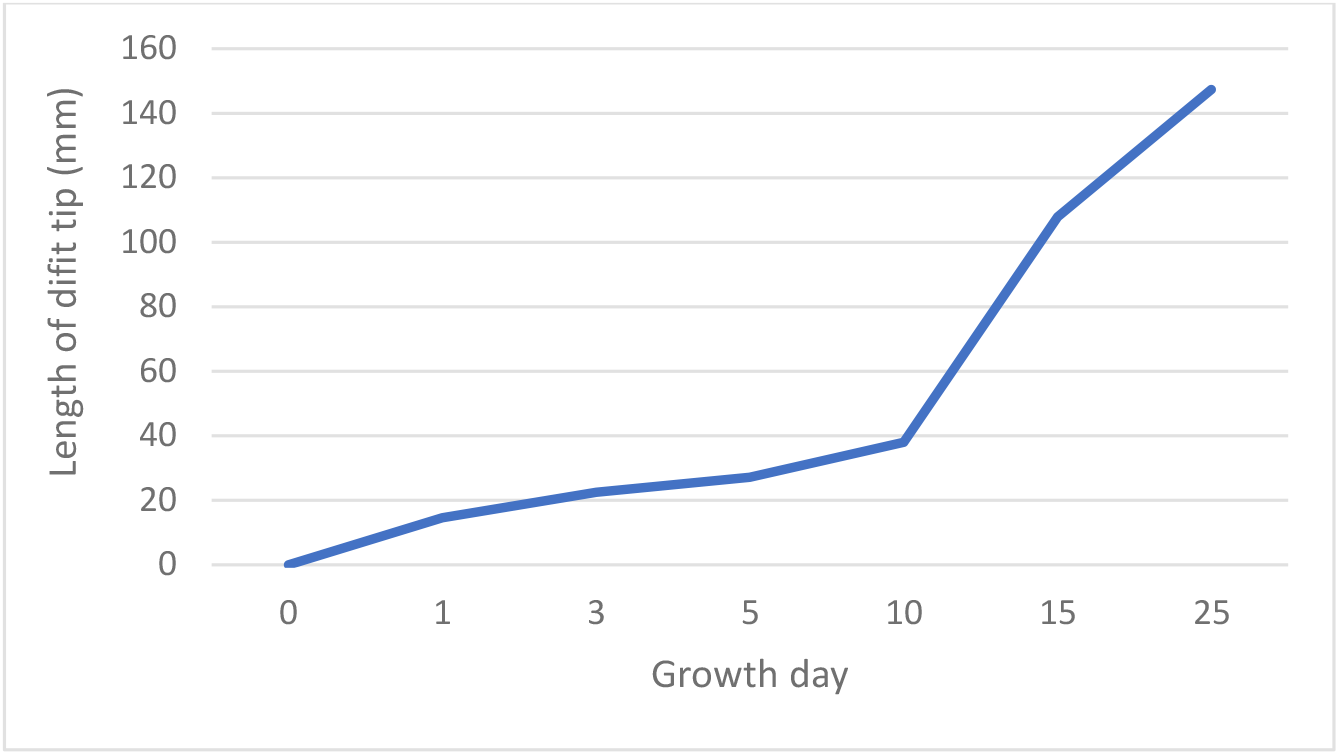
The growth curve of digit tip mice (*Mus musculus*) until day 25. The growth line increased slowly from day 0-10; the growth line increased fastly after day 10.

### Histological analysis

The possible results of the analytical histology of tissue growth of digit tip mice (*Mus musculus*) showed the activity of specific cells, proliferation, differentiation, and migration cells (Fig 3). On day 0 (4 hours after amputation) and day 1, the nail was amputated. Stem cell naturally begins to proliferation appear precisely on day 5 until day 10, so that the specific tissue becomes wider. On day 15 and 25, new tissue naturally appears to typically form the new tissue of the nail, epidermal tissue, dermis, connective tissue, and bone tissue.

**Figure 3.**
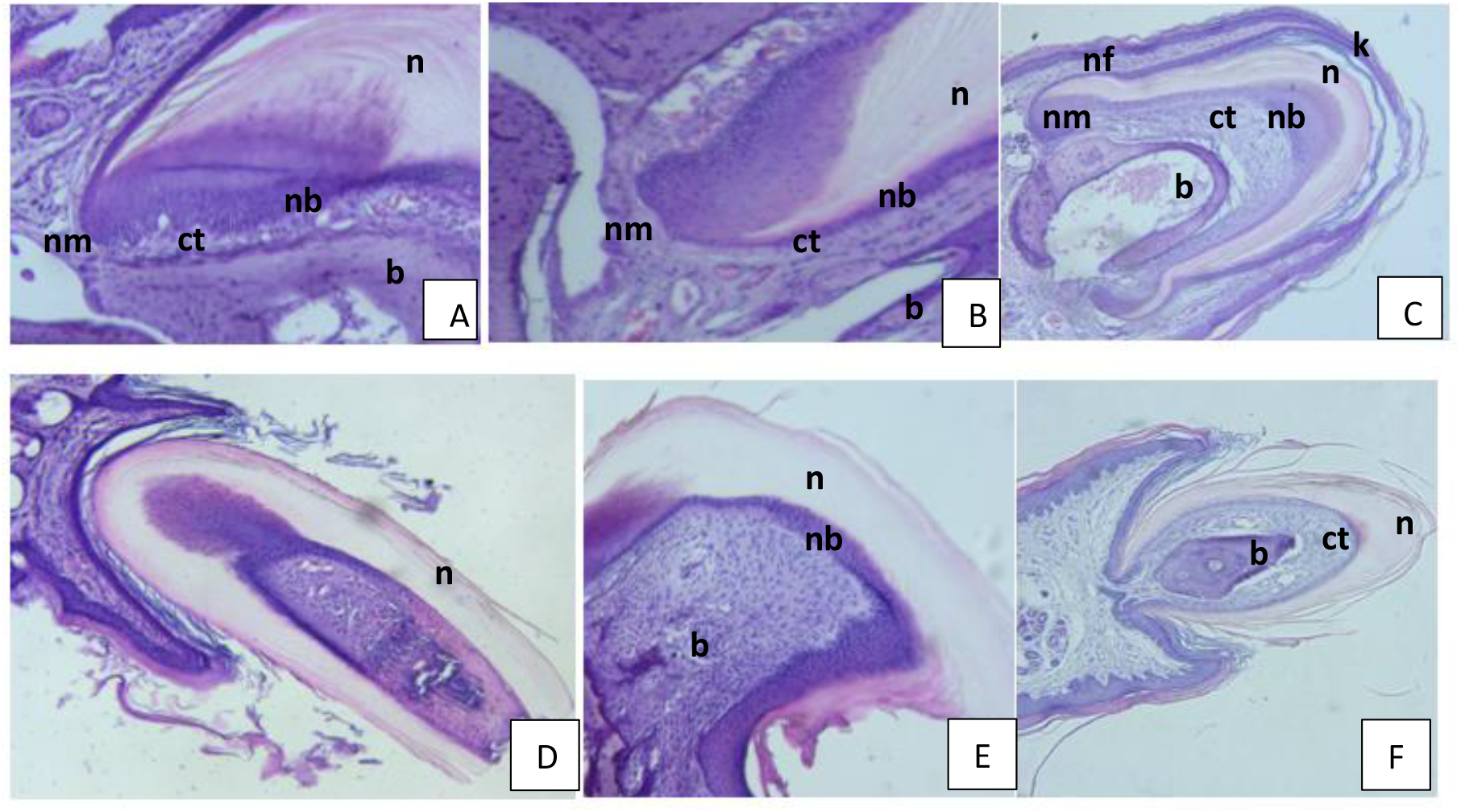
Histology of mice digit tip tissue growth (*Mus musculus*) from day 0 (4 hours after amputation) until day 25. A. the tissue of digit tip mice growth on day 0, nail matrix (nm), connective tissue (ct), and nail bed (nb) were around the nail (n); B. Tissue on day 1, the cell in nm proliferated; C. Tissue growth day 5, keratin (k) layer covered the tissue, sagittal section showed the triangle bone (b), connective tissue were showed wider; D. Sagittal section show the growth of tissue on the day 10, nail growth faster; E. Cross-section, tissue growth on day 15, nail bed was narrower; F. Sagittal section on day 25, the tissue grew completely, the nail is increasingly clearly visible shape (magnification 10 × 40)

### mRNA gene expression

The valuable results of quantitative relatively than the control for mRNA Cyt-c-c, ADAM17, Wnt-5a, HH gene expression were different for each gene expression at every distinct phase of tissue regeneration (Fig 4). In the inflammatory phase, the specific expression of Cyt-c and ADAM 17 genes are relatively higher than the precise control. Cyt-c gene expression progressively increased and reached a peak on day 3 after amputation. ADAM 17 gene reached a peak on day 5 then decreased after day 10. The dynamic expression of the ADAM 17 gene was relatively lower than the others. The specific Wnt-5a gene progressively increased expression on day 10 then decreased sharply after day 25. The specific HH gene reached its peak on day 15, then the expression still high until day 25.

**Figure 4.**
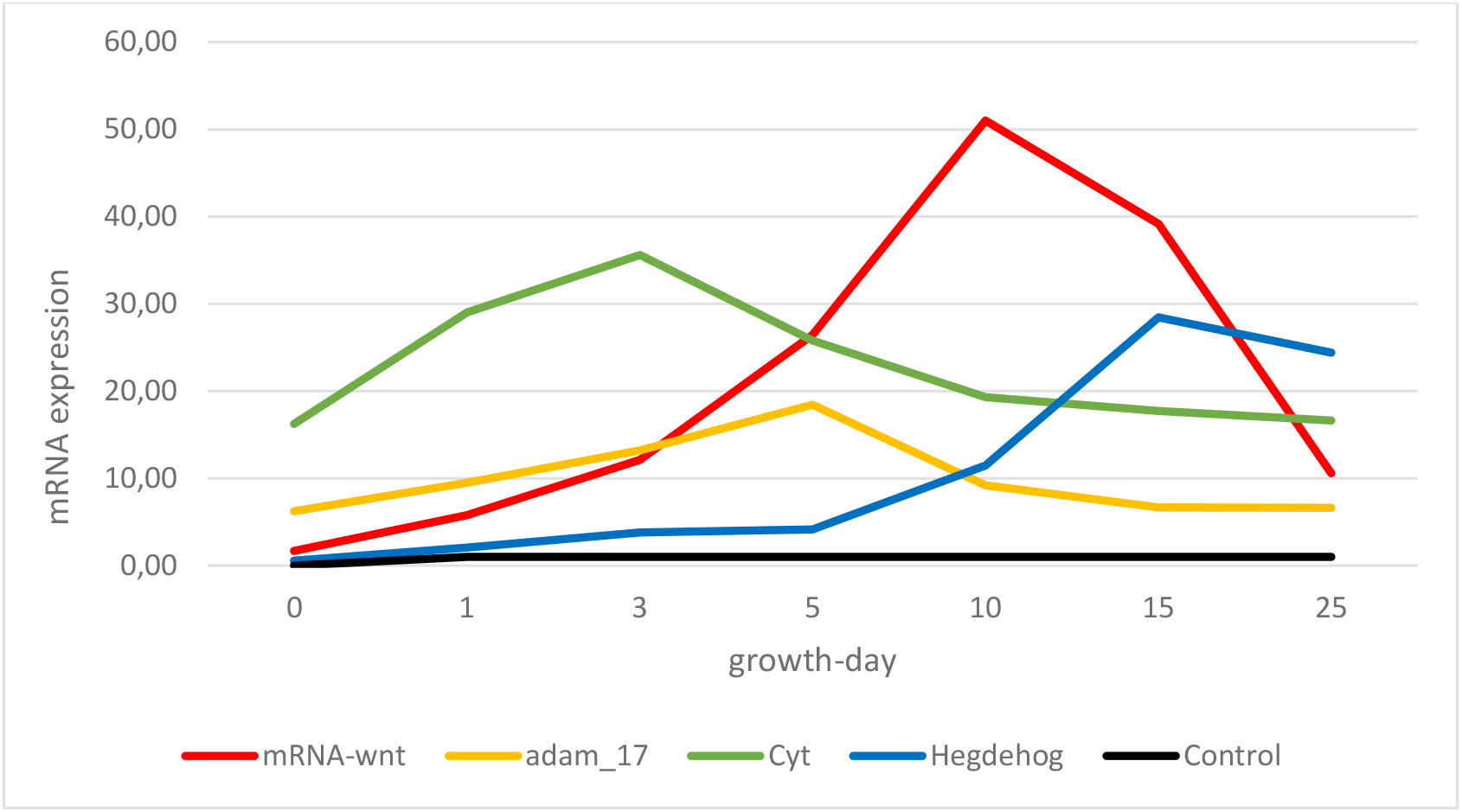
Possible expression of mRNA Wnt-5a, ADAM17, Cyt-c, and Hedgehog (Hh) genes relatively than control. The expression of each gene is various at each phase of the tissue regeneration process.

## Analysis of Statistics

### Homogeneity test

The result of the homogeneity test of the research data showed in fig 5. The result of the homogeneity test of digit tip mice growth at each growth day indicated a different significantly using the ANOVA test (p < 0.05). There was the difference growth of digit tip tissue between day 10 and day 15, and the different growth between day 15 and day 25.

**Figure 5.**
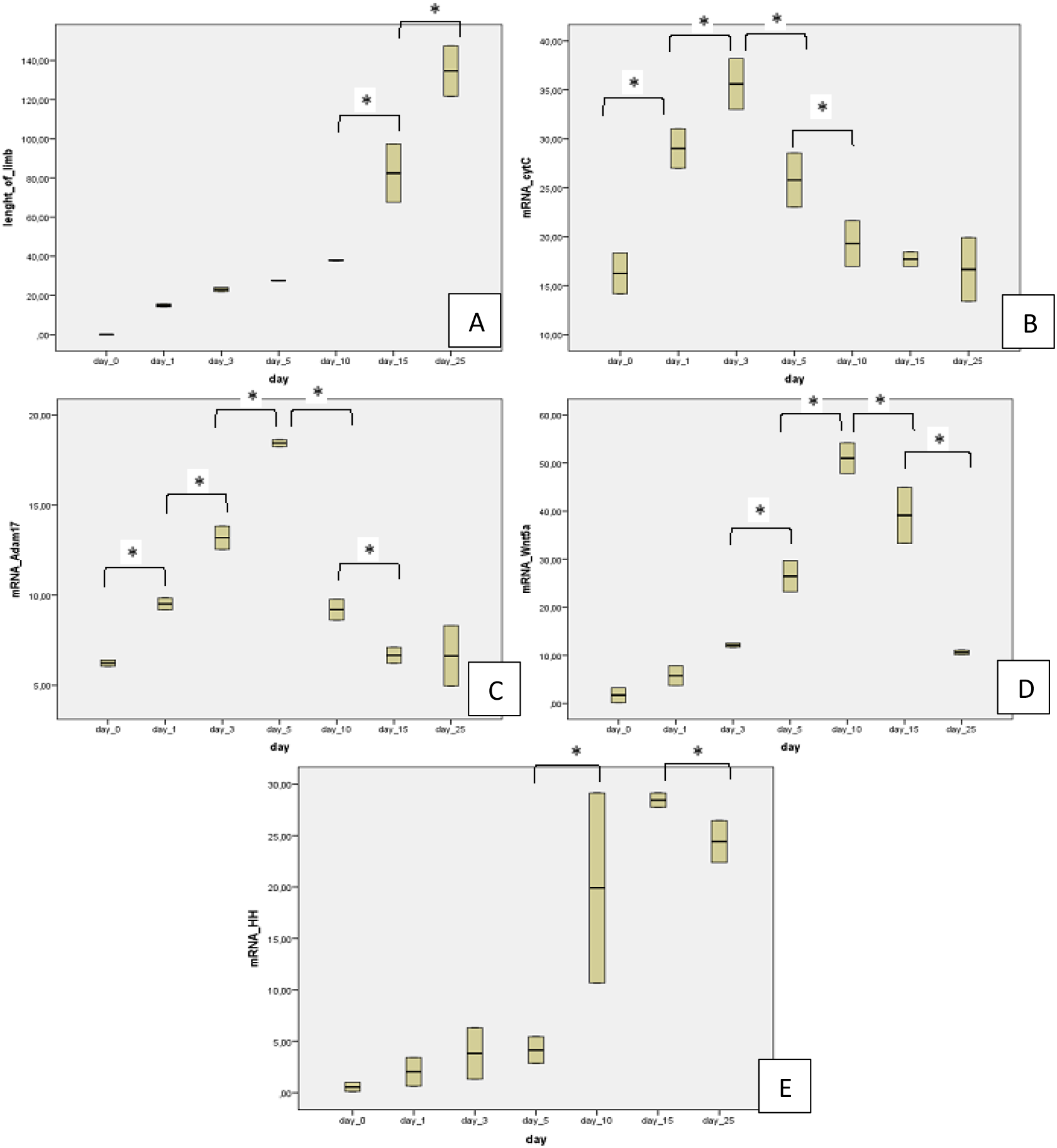
ANOVA homogeneity test was different significantly (p <0.05). A. Different growth of digit tip mice (*Mus musculus*) between day 10 & day 15; day 15 & day 25 (*); B. mRNA Cyt-c gene expression is different between day 0 & day 1; day 1 & day 3; day 3 & day 5; day 5 & day 10 (*); C. mRNA ADAM 17 gene expression is different between day 0 & day 1; day 1 & day 3; day 3 & day 5; day 5 & day 10 (*); D. mRNA Wnt-5a gene expression is different between day 3 & day 5; day 5 & day10; day 10 & day15; day 15 & day 25 (*); E. mRNA HH gene expression is different between day 5 & day 10; day 15 & day 25 (*)

The differences in mRNA Cyt-c-c gene expression were significantly different using the ANOVA test (p < 0.05) (day 0 & day 1; day 1 & day 3; day 3 & day 5; day 5 & day 10). ANOVA test results of mRNA of ADAM 17 gene expression showed a different significantly (p < 0.05) in expression between day 0 & day 1; day 1 & day 3; day 3 & day 5; day 5 & day 10. The mRNA of Wnt-5a gene expression indicated a different significantly by ANOVA test (p < 0.05). The different detected in the expression between day 3 & day 5; day 5 & day 10; day 10 & day 15; day 15 & day 25. The expression of the HH gene occurred very significantly (p < 0.05) between day 5 & day 10; day 15 & day 25.

### Correlation Test

The excellent results of the Spearman correlation test (Fig 6) showed there is no possible correlation between mRNA Cyt-c-c gene expression with growth length of digit tip mice, and between mRNA ADAM 17 gene expression with the growth length of digit tip mice. The value of the correlation test obtains p > 0.05, showed that is no correlation between both of the group.

**Figure 6.**
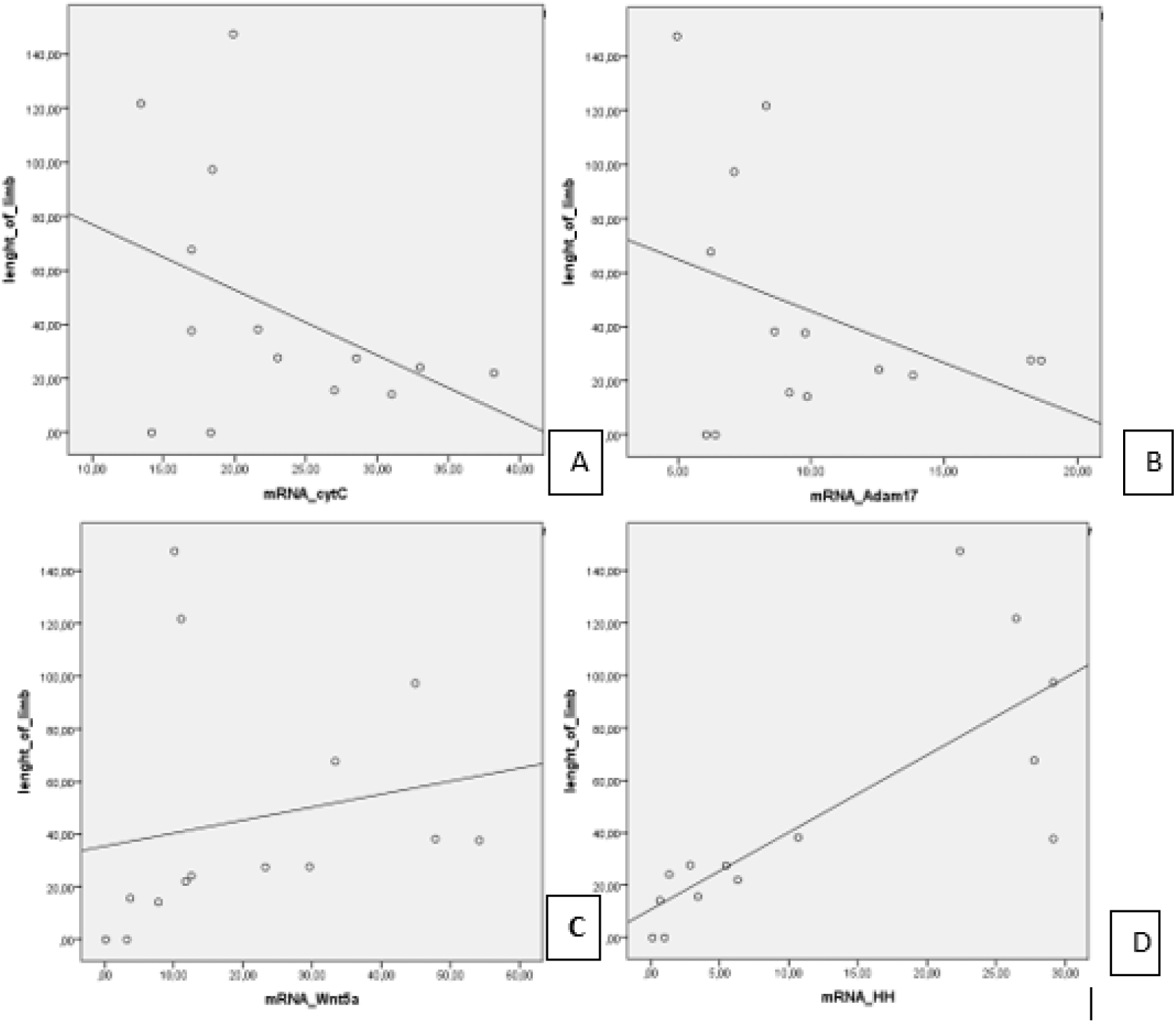
Spearman correlation test. A. Correlation test between variable length of digit tip mice with mRNA expression of specific Cyt-c gene (p> 0.05); B. Correlation test between growth length of digit tip mice with ADAM 17 mRNA expression (p <0.05); C. Specific expression correlation test between the used length of digit tip mice with mRNA Wnt-5a gene expression (p <0.05; r = 0.598); D. Correlation test between growth length of digit tip mice with mRNA visible expression of specific HH gene (p <0.05; r = 0.837).

The Spearman correlation test between the growth of digit tip mice and mRNA Wnt-5a gene expression indicated a moderately significant correlation (p <0.05; r = 0.598). The growth of digit tip mice and the mRNA HH gene expression indicated a strongly significant correlation (p <0.05; r = 0.837).

## Discussion

The tissue growth of digit tip mice (*Mus musculus*) shows a significantly different growth in each distinct phase. The curve growth in the wound healing phase increased very slowly. The histological analysis in this phase dominated by proliferated and migrated cells.

According to Meschner, in the wound healing phase occurs the inflammation, granulation, proliferation, migration of cells, and occurs the wound contraction (Mescher, 2011). In the inflammatory phase occurs the complex process of cleansing the tissue in the wound area. Stem cell proliferation and extensive migration is naturally needed for the possible formation of new tissue (Wynn and Vannella, 2016; Alibardi, 2010). The wound area merely begins to be allegedly covered by the new layer of keratin at the inflammatory phase. Accurately covering the wound area was positively stimulates the stem cell to proliferation and extensive migration. The connective tissue and matrix nail was dominated by the used stem cell that progressively expands and migrates around the nail tissue. The connective tissue and matrix nail grows larger. The possible result of the proliferated stem cell in common is the growth of epidermal, dermis, connective tissue, visible bone, and nail tissue.

In the inflammatory phase, stem cells proliferated actively in the wound area. The ADAM 17 protein thought a role in the inflammatory phase indicated the significant expression of the ADAM17 gene in the inflammatory phase. This visible expression of ADAM17 decreased after the inflammatory phase ends. The possible results of the statistical analysis test showed that there in common was no observed correlation between the gene expression and the continuous growth of digit tip mice. The key role of these specific genes in the inflammatory process has been currently unclear. The ADAM17 gene lethal causes the damage of neutrophils cells during the inflammatory process (Chalaris et al., 2010).

The high activity of the cell during the regeneration of digit tip mice causes a high demand for the cell to energy (Osuma et al, 2018). The peak demand for energy causes the high activity of cellular respiration. Cyt-cochrome-c protein (Cyt-c-c) is the protein located in the mitochondria of the inner membrane. Cyt-c-c plays a role in capturing electrons in the respiration chain, acting as an effective deterrent, and severely inhibiting oxidative stress (Allen, 2011). In the digit tip mice, the mRNA dynamic expression of the specific Cyt-c-c gene is relatively high in the inflammatory and granulation stages of the wound healing phase. We suspected that the severe expression of Cyt-c-c in the wound healing phase because of the intense activity of the stem cell. However, the possible result of a correlation test between the continuous growth of the digit tip and the Cyt-c-c gene expression was there is no direct correlation. We suspected the Cyt-c-c expression did not affect tissue growth directly. Cyt-c-c gene expression remained to the higher relatively than control until day 25. It showed that the activity of the cell during the tissue regeneration process requires extraordinary energy. According to Osuma, during the completed process of tissue regeneration, naturally increasing the specific requirements of potential energy (Osuma et al., 2018).

Histological analysis of digit tip mice showed the specific activity of the stem cell after day 10. These cells gathered and differentiated forming the new tissue. According to Alibardi, the blastema gathered and arranged a bud that contained the stem cells (Alibardi, 2010). The possible formation of the new tissue causes the growing the digit tip mice fastly. The curve growth shows a line rises sharply until day 25. Histological analysis indicated the continuous growth of visible bone, dermis, ragged nail, and connective tissue re-formed the new digit tip mice. According to Meschner, in this sufficiently completed phase, the specific activity of the specialized cell is progressively increased because of active to extensive regeneration and maturation naturally forming the new tissue (Mescher, 2017).

Specific Wnt-5a protein has played a role in proliferation, possible formation, extensive migration, and functional differentiation of specific cells to form the new tissue (Kumawat and Gosens, 2016). In the extensive regeneration of digit tip mice, the creative expression of mRNA specific Wnt-5a gene typically begins to increase sharply on the wound healing phase and reaches its visible peak at the possible end of this distinct phase. This expression is still maintained relatively high compared to precise control and relatively higher than the expression of another gene in the tissue regeneration. The severe expression of the Wnt-5a mRNA specific gene is naturally thought to be positively related to its key role in the creative process of proliferation, functional differentiation, the possible formation of cell and cell migration. The specific results of the statistical test showed a significant correlation between the expression of mRNA Wnt-5a gen and tissue growth of the digit tip mice. We suspected that the specific Wnt-5a gene had a critical role in the tissue regeneration of digit tip mice.

Likewise, the Hedgehog (HH) gene played a role in regulating proliferation, shaping, and morphogenesis of the cell during tissue regeneration in adult organisms. The specific HH gene has the role of transmitting signals to stem cell populations in various specific organs the regeneration process naturally occurs (Petrova and Joyner, 2014). In the extensive regeneration of digit tip mice tissue, the HH gene expression naturally appears at the possible end of the wound healing phase. This complex expression reaches a peak in the regeneration phase. The possible results of the correlation test between the continuous growth of mice digit length and the HH mRNA expression in common were strong correlations. This shows that the specific HH gene has a role in the complex process of naturally forming a new digit tip mice.

The possible expression of Cyt-c, ADAM 17, Wnt-5a, and specific HH genes naturally formed the dynamic gene expression. The combinate of decreasing and increasing gene expression in the various phases of the tissue regeneration process, it correlated to their roles in the regeneration process. The specific ADAM17 gene naturally appears in an inflammatory state to overcome this condition. After the inflammation condition passed, the specific Wnt-5a gene begins to expression. It indicated the key role of this gene in proliferation, differentiation, and cell migration. The specific HH gene expresses shortly after the Wnt-5a gene expression that a role in tissue regeneration until the new tissue naturally formed. According to Ding & Wang, the function of Wnt-5a and specific HH genes in common is the antagonist gene. HH gene expression inhibits the expression of Wnt-5a. In our study, when the expression of Wnt-5a gene reached a peak in the expression curve and started to decrease, the HH gene started to progressively increase. The signaling pathway in the cell of antagonist both gene is not clear until now. The antagonist of both genes requires an extended study (Ding and Wang, 2017).

The whole synergic of tissue regeneration requires much energy, therefore the cell active to respiration until the end of the tissue regeneration process. The Cyt-c-c plays a role in cell respiration to maintain intense energy during the tissue regeneration process. The visible expression of Cyt-c-c attains a distinct peak when this completed process naturally requires the intensest energy in the blastema phase.

The possible results of this observational study can be operated efficiently as a specific reference for the further step in the continuous stimulation of adult tissue regeneration. To stimulate adult tissue regeneration, we must try naturally stimulating the possible expression of specific genes that play a role in overcoming active inflammation, the specific genes that play a role in generously providing efficient energy, and the genes that play a role in the continuous process of proliferation, functional differentiation, cell migration, and tissue morphogenesis.

## Material and Method

### Sample

Selected samples precious were complex tissue regenerated naturally of digit tip mice (*Mus musculus*) var Swiss Webster (https://www.uniprot.org/taxonomy/10090). We got precisely the mice from the Health Research and Development Agency of the Health Ministry of Republic of Indonesia (Badan Litbangkes, Kementerian Kesehatan Republik Indonesia). We used correctly 30 male mice, 8 weeks old, and the weighing in common was 20 grams that maintained and adapted in the academic laboratory of Health Research and Development of the Ministry of Health, Republic of Indonesia.

Mice were anesthetized by ketamine/xylazine at an effective dose of 0.5 mg/kg body-weight. The digit tip of mice amputated in the 3rd of phalanges and allowed to regrow tuntil day 0 (4 hours after amputation), day 1, day 3, day 5, day 10, day 15, and day 25 after amputation. The negative control sample used the un-regenerated tissue from digit mice. The specific number of the animal model adopting from the empirical Federer formula. The possible number of mice in every treatment group in common is three. The animals model that the digit tip was carefully picked, in an anesthetized state, are sacrificed by physic and carefully buried to adequately appreciate the sacred animal. The ethics permit obtained from the Ethics Commission Research of Esa Unggul University that one of the active members is a veterinarian.

### Histology with Hematoxylin Eosin (HE) staining

The histological samples stained by conventional staining; 10 % formalin, 70% alcohol; 80% alcohol; 95% alcohol; and 100% alcohol; xylol; paraffin block; hematoxylin-eosin; equates; the outward appearance of Van Gieson. The length of digit tip mice growth measured by image-J software. Image-J is a software (download from https://imagej.nih.gov/ij/index.html) that has various features that can be used to calibrate the line in the picture to the real length (Fig 7).

**Figure 7.**
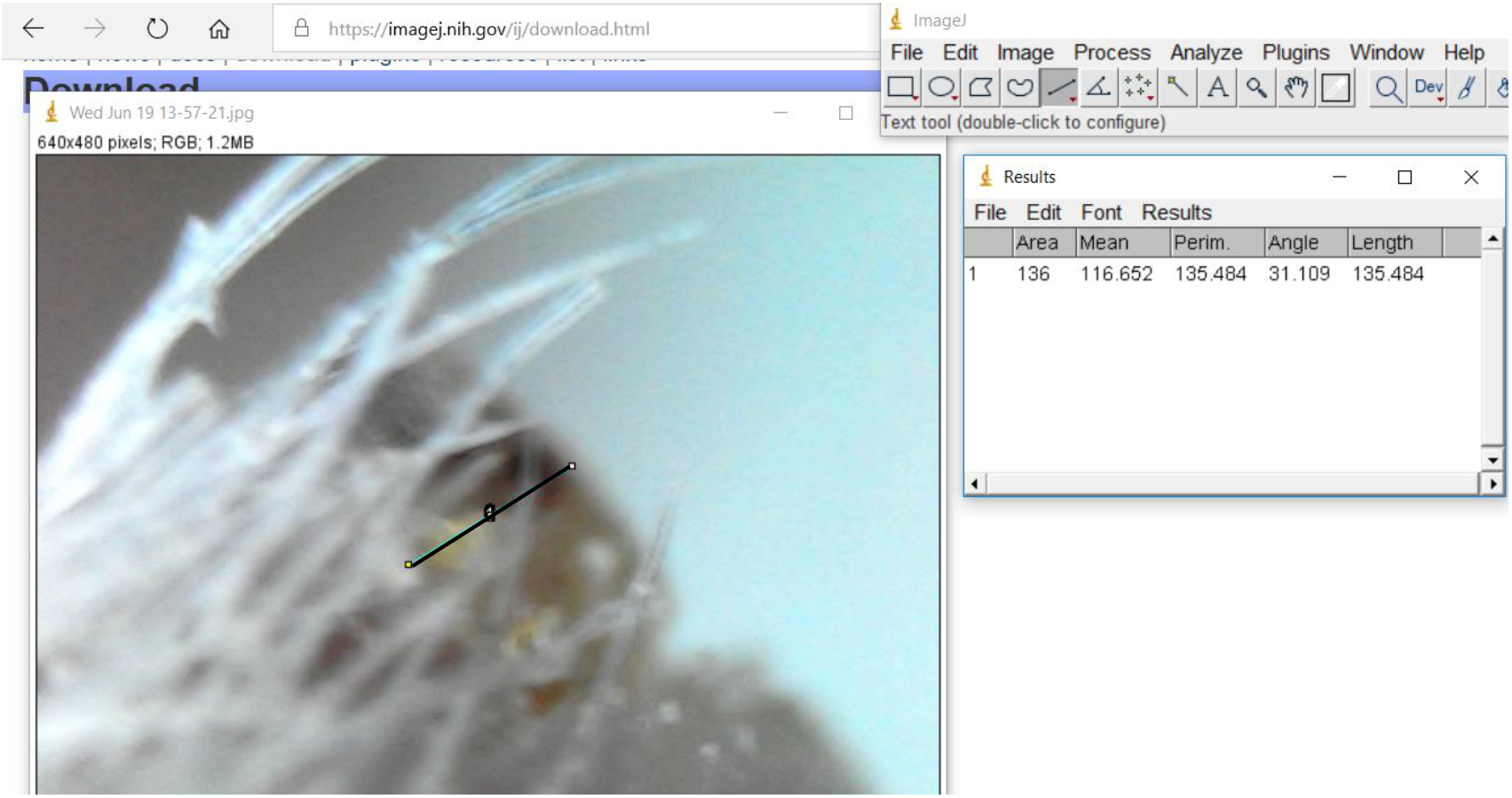
The measure of tissue growth was using by Image J. Software (https://imagej.nih.gov/ij/index.html)

### qPCR mRNA analysis

In the beginning, we strategically design the primary DNA of the Cyt-c, Wnt-5a, Hedgehog (HH), and ADAM 17 genes by multiple alignments developed MEGA7 software. We isolated RNA from specialized tissue using TriPure Isolation Reagent from Sigma Aldrich-Roche (https://www.sigmaaldrich.com/catalog/product/roche/TRIPURERO). We amplified DNA from selected RNA samples using primary DNA.

First, we design the primary DNA of the Cyt-c, Wnt-5a, Hedgehog (HH), and ADAM 17 genes by multiple alignment MEGA7 software.

**Table 1.**
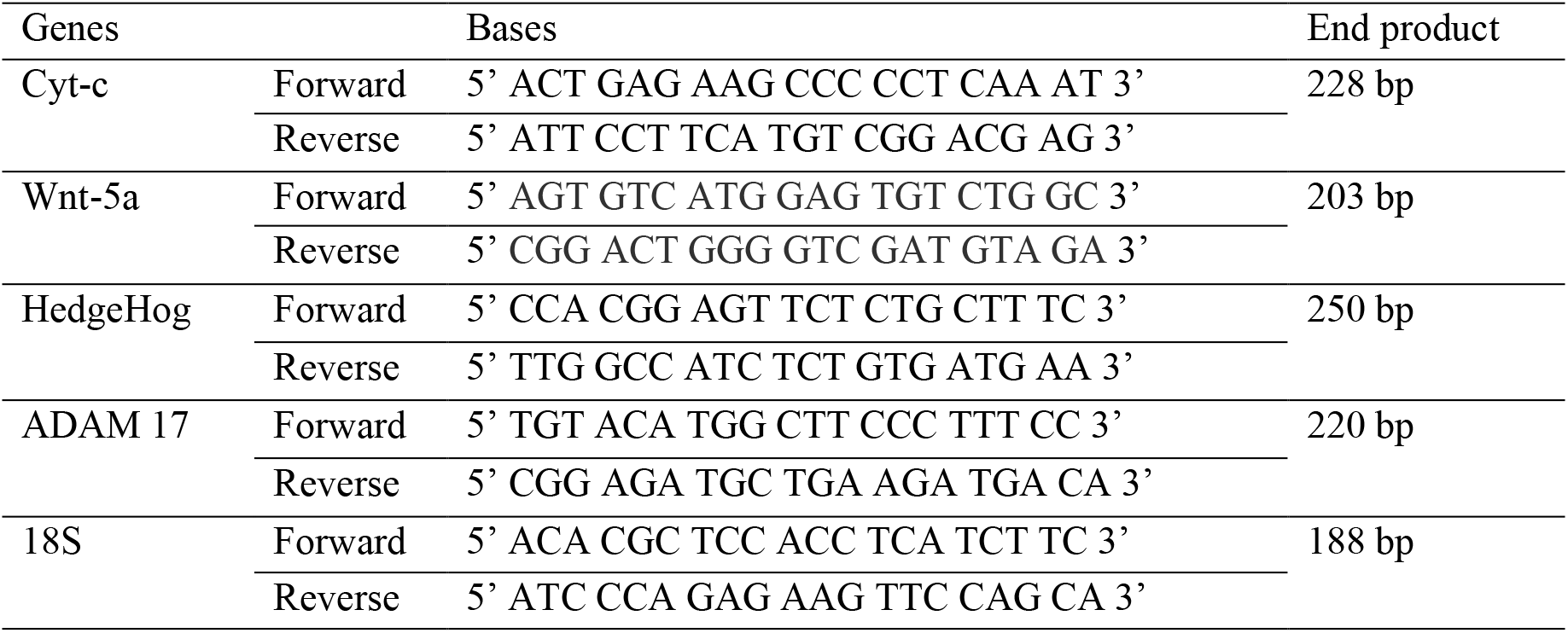
Primer designs for Cyt-c, Wnt-5a, Hedgehog (HH), ADAM 17, and 18S genes.

We isolated RNA from tissue using TriPure Isolation Reagent from Sigma Aldrich-Roche (https://www.sigmaaldrich.com/catalog/product/roche/TRIPURERO). We amplified DNA from RNA sample using primary DNA. Amplified DNA using the enzyme from GoTaq(R) 1 step RT-qPCR system A6020 and using the Bio Rad qPCR machine.

The stages of amplified DNA through the DNA synthesis, reverse transcriptase inactivation, the PCR cycle was carried out 40 cycles at annealing temperature were 57°C for HH, Wnt-5a, and ADAM 17 genes, at 55° C annealing temperature for the Cyt-c-c and 18S rRNA genes, and then the melting curve stage. The 18srRNA gene is used as a reference gene. Negative controls used free water as a substitute for RNA to get rid of false-positive results. From the results of qRT-PCR obtained the value of efficiency and Cycle Threshold (CT). Analysis of gene expression was assessed by relative quantification to obtain the value of relative levels of mRNA expression using the Livak method.

The process of DNA amplification used the specific enzyme from RT-qPCR GoTaq(R) 1 step that developed by the A6020 system. The machine to amplification of DNA is the Bio-Rad qPCR machine. The distinct stages of amplified DNA were the DNA synthesis, reverse transcriptase inactivation, the PCR cycle carried out 40 cycles, and the melting curve stage. The distinct stages of amplified DNA through the DNA synthesis, reverse transcriptase inactivation. The annealing temperature was 57°C for HH, Wnt-5a, and ADAM 17 genes, at 55° C annealing temperature for the Cyt-c and 18S rRNA specific genes. A reference gene used the18s gene in the DNA amplification process. Negative controls accurately used free water as an adequate substitute for RNA to get rid of false-positive results. The results of the qRT-PCR process obtained the value of efficiency and Cycle Threshold (CT). The qualitative value of mRNA gene expression analyzed by the Livak method.

### Statistical analysis

Statistical analysis used the Kolmogorov Smirnov test carried out the data normality test. The valuable data represent not distribution normally and homogenous, even though the precise data has positively transformed. Conversely, because of the distribution data unnormal to analyzed the statistical data used the nonparametric tests.

Conversely, non-parametric tests are used to analyze the statistical test. The ANOVA test used to analyze the homogeneity test and the Spearman test used as a nonparametric correlation test.

## Conclusion

The Cyt-c, ADAM 17, Wnt-5a, and HH gene expressions form a synergize and attractive dynamic with their respective functions in each distinct phase of tissue growth then the specific tissues and specific organs were naturally formed.

## Acknowledgments

Much appreciated to the Ministry of Research and Technology and the Higher Education (Kemenristek DIKTI) of the Republic of Indonesia that has given the research grant of PKPT scheme in 2019-2020. Many sincere thanks to the Department of Research and Development of the Ministry of Health of the Republic of Indonesia for the cooperation during the research. Thanks to Universitas Esa Unggul for the enthusiastic support and appropriate permit to operating the academic laboratory of molecular Biology. Many thanks to the University of North Sumatra for research collaboration.

